# Ultra-Low Colcemid Doses Induce Microtubule Dysfunction as Revealed by Super-Resolution Microscopy

**DOI:** 10.1101/2020.08.13.249664

**Authors:** Ashley M Rozario, Sam Duwé, Cade Elliott, Riley B Hargreaves, Peter Dedecker, Donna R Whelan, Toby D M Bell

## Abstract

Microtubule-interacting drugs, sometimes referred to as antimitotics, are used in cancer therapy to target and disrupt microtubules. However, their side effects require the development of safer drug regimens that still retain clinical efficacy. Currently, many questions remain regarding microtubule-interacting drugs at clinically relevant and ultra-low doses. Here, we use super-resolution microscopies (single molecule localization and optical fluctuation based) to reveal the initial microtubule dysfunctions caused by nanomolar concentrations of colcemid. Short exposure to 30 - 80 nM colcemid results in aberrant microtubule curvature while microtubule fragmentation is detected upon treatment with ≥100 nM colcemid. Remarkably, even ultra-low doses (5 hours at <20 nM) led to subtle but significant microtubule architecture remodeling and suppression of microtubule dynamics. These challenges to microtubule function represent less severe precursor perturbations compared to the established antimitotic effects of microtubule-interacting drugs, and therefore offer potential for improved understanding and design of anti-cancer agents.

## Introduction

Microtubules (MTs) form part of the cytoskeleton and have many essential roles in the cell, including maintaining cell shape and supporting transport of organelles and vesicles^1^. To fulfil these roles, MTs form intracellular networks comprising 25-nm wide hollow filaments, polymerized from *αβ* -tubulin dimers. Additionally, MT dynamics through tubulin assembly and disassembly contribute forces to segregate chromosomes during mitosis^2,3^. These critical functions make MTs vulnerable to bacterial pathogens^4^ and have recently been shown to be susceptible to viral subversion^5,6^. The importance of MT function also presents as a viable target for cancer therapy^7,8^ and some compounds approved as anticancer drugs (Paclitaxol^9^ and Vinblastine^10^) are known to bind tubulin and alter MT filament stability. Colchicine, a naturally occurring compound^11^, induces MT depolymerization and is used to treat inflammatory diseases including gout and familial Mediterranean fever^12^. The use of colchicine, however, is limited by its toxicity and poorly defined dosing thresholds between non-toxic, toxic and lethal^13–15^. It can cause neurotoxicity^16^ and is associated with renal and liver failure in cases of colchicine poisoning^13^. Despite these outcomes, colchicine still presents some potential as an anticancer compound where low concentration doses reduce proliferation of cholangiocarcinoma cell lines, as well as decrease tumour size in mouse models^17^. Several synthetic derivatives of colchicine are being clinically tested and show promise to treat cancers in the future^18^.

The physiological outcomes of treatment with MT-interacting drugs vary between patients, making it difficult to standardise a therapeutic dose. This complication can be due to the individual’s resistance toward the drug, manifested either by cancer-driven genetic changes or acquired resistance over several drug treatments^19^. The onset of adverse outcomes may also be due to the drug’s non-specificity for cancer cells, meaning non-cancerous cells also suffer some effects. MT-interacting drugs are well-established as being able to arrest cells in mitosis, preventing normal division and potentiating apoptosis. Antimitotic effects also include mitotic structural aberrations such as the formation of multiple spindles and defective daughter cells with unnatural amounts of genetic information^20^. However, an emerging hypothesis explaining the action of MT-interacting drugs is that non-mitotic effects are key to their efficacy^21–25^. This is increasingly important for understanding therapeutic mechanisms because these drugs bind tubulin regardless of cellular phase. As a result, MT structure and dynamics become altered, affecting MT-dependent functions including intracellular transport, signalling cascades and cell motility. This may also trigger the dysfunction of MT-associated organelles such as mitochondria^26^ and actin^27^. Given the established importance of MTs in everyday cell biology, any form of MT dysfunction will undoubtedly have significant outcomes on cell health and by extension, patient health. Ideally, chemotherapeutic MT-interacting drugs should be used at the lowest dose possible that provides anticancer results while minimizing off-target damage. Historically, these drugs have been studied predominantly for their antimitotic outcomes (mitotic arrest, daughter cell mutations). However, these effects are typically lethal for cells and better represent high level toxicities from high drug doses. It is increasingly clear that the much lower doses relevant in clinical use may not induce significant mitotic effects and that their mechanism is instead to affect non-mitotic MT function. These are inherently more subtle than antimitotic effects and have proven difficult to characterize using conventional imaing and biochemistry. Therefore, we set out to better understand clinically relevant mechanisms of MT-interacting drugs using complementary super-resolution microscopies which enable both sub-diffraction and dynamic live-cell observations of MT structure and function.

These super-resolution techniques build on fluorescence microscopy which has long been a cornerstone tool for visualizing intracellular features, albeit with limited imaging resolution of ∼200-300 nm due to the diffraction of light. In order to observe the sub-diffraction architecture of the MT network, super-resolution methods are necessary^28^. Single molecule localization (SML) approaches such as *d*STORM (direct stochastic optical reconstruction microscopy) achieve as good as 20 nm resolution, imparting a 10-fold improvement over the conventional limit of optical microscopy^29^. While SML achieves excellent resolution gain it is not well-suited to live-cell dynamic imaging because of the typical need for long acquisition times and high laser powers. Super-resolution optical fluctuation imaging (SOFI)^30,31^ has proven much more compatible with live-cell microscopy because of comparatively mild preparation and acquisition conditions. SOFI achieves sub-diffraction resolutions through a post-processing approach, using statistical analysis of the fluorescence dynamics of the fluorophores. Like *d*STORM, SOFI can be conveniently performed on conventional wide-field microscopes with EM-CCD or sCMOS detectors. SOFI employs reversibly switching fluorescent proteins (RSFPs) such as Dronpa^32^ that photoswitch in response to low laser excitation (tens of mW/cm^2^) and entails much shorter acquisition periods (seconds to tens of seconds) for individual frame generation. Though the resolution gain is, in principle, unlimited, the realities of RSFP labelling density, switching kinetics and photostability have so far restricted practical efficiency of SOFI to an overall 4-fold resolution improvement at best^33^. Used in parallel, *d*STORM and SOFI complement each other providing super-resolution detail and live-cell relevance and context, respectively, for a holistic perspective of the biological system in question.

Here, we applied *d*STORM and SOFI to visualize initial MT perturbations caused by short exposure to low concentrations of colcemid. Colcemid is a derivative of colchicine commonly used in laboratory settings at higher doses to arrest cells in metaphase for karyotyping assays^34,35^. We detected significantly altered filament curvature upon 5 hour treatments with colcemid concentrations as low as 7 nM, with visibly appreciable defects present in cells dosed with 50 nM, while more pronounced filament curvature alterations were detectable at 80 nM. Interestingly, this correlates with the maximum concentration detected in blood plasma of patients treated with a therapeutic dose of colchicine^36^. Measured filament curvatures were as high as 2 rad/*µ*m in 50-80 nM treated cells, a value which previous studies associated with MT breakage^37,38^. Increasing colcemid concentration to 100 nM and 200 nM resulted in less abundant cellular MT filaments which appeared shorter, demonstrating that filaments had become fragmented. To further probe the effects of ultra-low colcemid doses (<30 nM), we applied a SOFI time-lapse approach to capture dynamics of individual filaments with a temporal resolution of 20 s (one SOFI image every 20 s) for up to 9 minutes. Compared with untreated cells, as low as 18 nM colcemid caused individual filaments to be significantly less active, both in growth and shrinkage, indicating suppression of MT dynamics. Despite a general consensus that hindering filament dynamics is a mechanism for therapy, direct observation and quantification of filament activity in living cells remains challenging, especially at clinically relevant ultra-low doses. The super-resolution assays developed here have revealed new insights into the effects of the MT-interacting drug colcemid, highlighting the potential therapeutic role of induced suppression of dynamics and aberrant curvature. This study also establishes a new standard for probing the sub-diffraction landscape of MT network perturbation which will be key in improved understanding, and further development, of MT-interacting drugs.

## Results

### Low doses of colcemid cause remodelling of microtubule architecture

*d*STORM was used to detect sub-diffraction effects of low doses of colcemid by immunolabelling tubulin in fixed HeLa cells treated with 0 - 200 nM colcemid for 5 hours (Figure 1). Visual inspection of the resulting images of whole cell MT architecture reveals clear changes to filament shape, abundance and arrangement in the cells treated with higher concentrations. However at 7 nM and 30 nM colcemid, MTs appear by eye to be similar the control, having relatively linear filaments oriented toward the cell’s edge from the centre. At 50 nM colcemid, while filaments are still mostly linear, some filament sections have become visibly more curved. 65 nM and 80 nM colcemid treatment resulted in more pronounced changes to filament curvature throughout the cell. Treatments with 100 nM and 200 nM colcemid results in the most striking effects, including a lower number of filaments that are relatively shorter, prescriptive of the well-established MT depolymerizing ability of colcemid (and colchicine)^39^. We hypothesize that these filament fragments are likely also derived from longer filaments that have broken due to excessive curvature. There is also an increase in “speckled” signals dispersed throughout the cytoplasm that could be MT filament constituents (*αβ* -tubulin dimers/oligomers). Some cells treated with 200 nM showed a remaining small dense structure, likely the MT organising centre (MTOC), located adjacent to the nucleus (SI Figure 1, bottom row, yellow arrows). This is consistent with the current understanding that MT depolymerization is induced from the periphery end (+ end) of filaments^40^.

**Figure 1.**
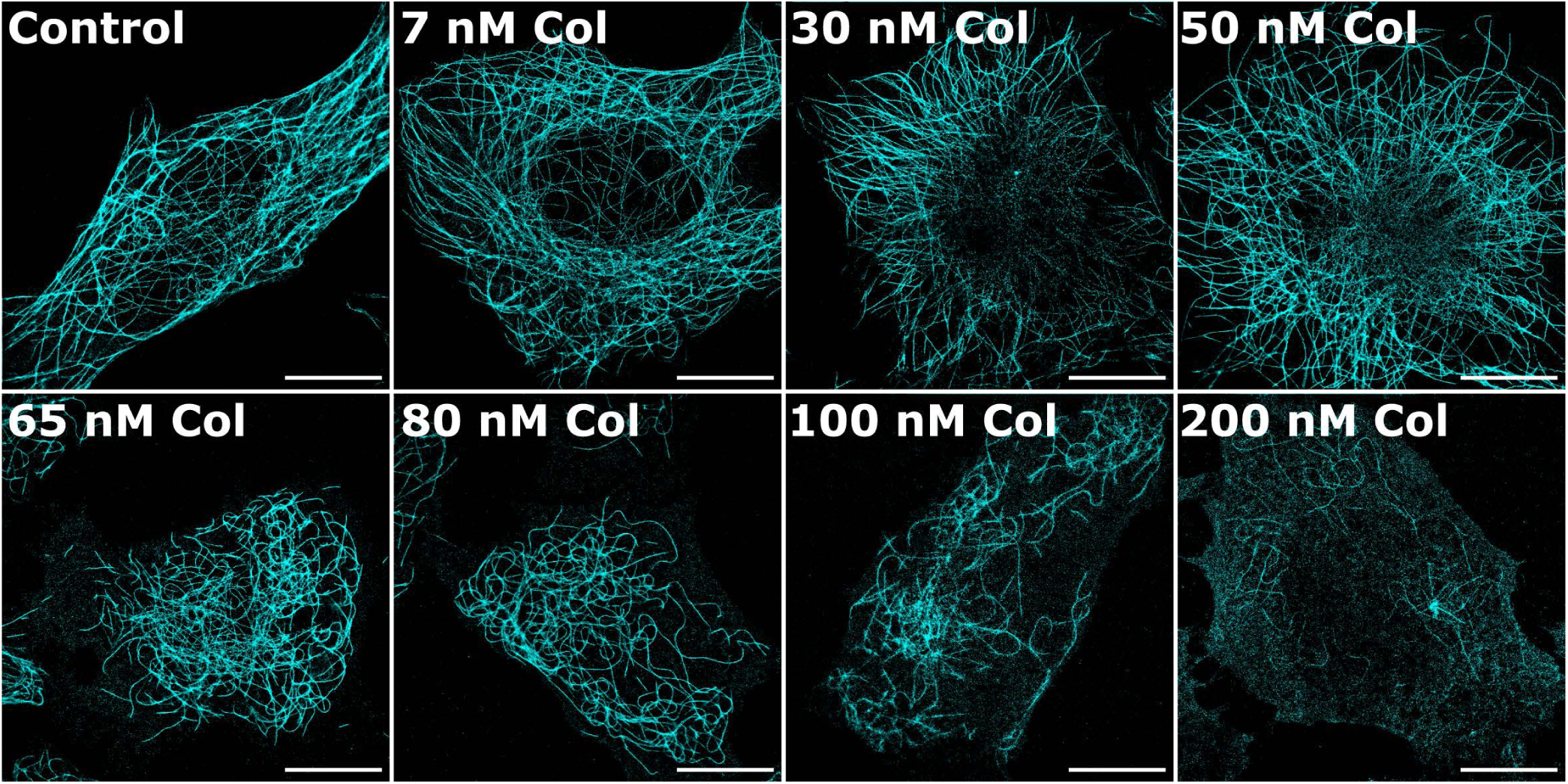
Single molecule super-resolution imaging (*d*STORM) of MTs in HeLa cells treated with colcemid reveals increasingly aberrant filament curvature. Cells were treated with colcemid at the specific concentrations for 5 hours before fixation. Cells were immunolabelled for tubulin with Alexa Fluor 647, imaged and super-resolution images rendered using rapi*d*STORM. Increasingly abnormal curvatures and fragmentation of the MT architecture can be visually detected with very significant losses observed at 100-200 nM. Images are representative for each treatment. N > 20 cells per condition. Scale bars = 10 *µ*m.

Notably 80 nM colcemid-treated cells wherein aberrant filament curvature could be detected by visual appraisal, coincides with the maximum concentration found in blood plasma of patients treated with therapeutic doses of colchicine^36^. Blood concentrations above this, and more comparable to the 100 - 200 nM doses which we show cause gross MT network destabilization, have been associated with lethal doses^41,42^. In these studies, drug concentrations were measured from solution in contact with patient’s cells, similar to our experiments where colcemid concentration was inferred from the final growth medium concentration. This allows for closer comparison between effects from single cell experiments and therapeutic doses, without measuring intracellular drug concentrations that may vary due to cellular uptake and metabolism.

To quantify filament curvature from *d*STORM images, we used SIFNE (SMLM image filament network extractor)^43^ to calculate the extent of curvature at each pixel along a traced filament. Figure 2a compares representative filaments from *d*STORM images of control and 80 nM colcemid cells with the same filaments traced by SIFNE and coloured for curvature between 0 and 2 rad/*µ*m. Each curvature value is obtained as the reciprocal of radius of a circle derived from the rate of curvature (see SI Figure 2a) and determined every 20 nm along a continuous filament. The distribution of curvature values from each cell were then histogrammed with bins of 0.1 rad/*µ*m (∼5.7°). Figure 2b shows curvature distributions from both a cell with mostly linear filaments (control) and one with more curved filaments (80 nM colcemid treatment). Each cell analysed provided at least 14,000 curvature values from several hundred microns of summed filament length. Curvatures at ∼1 rad/*µ*m relate to intermediate filament curves whereas those beyond 1.5 rad/*µ*m are associated with more extensive filament curvatures that sometimes exhibit 180° loops^43^.

**Figure 2.**
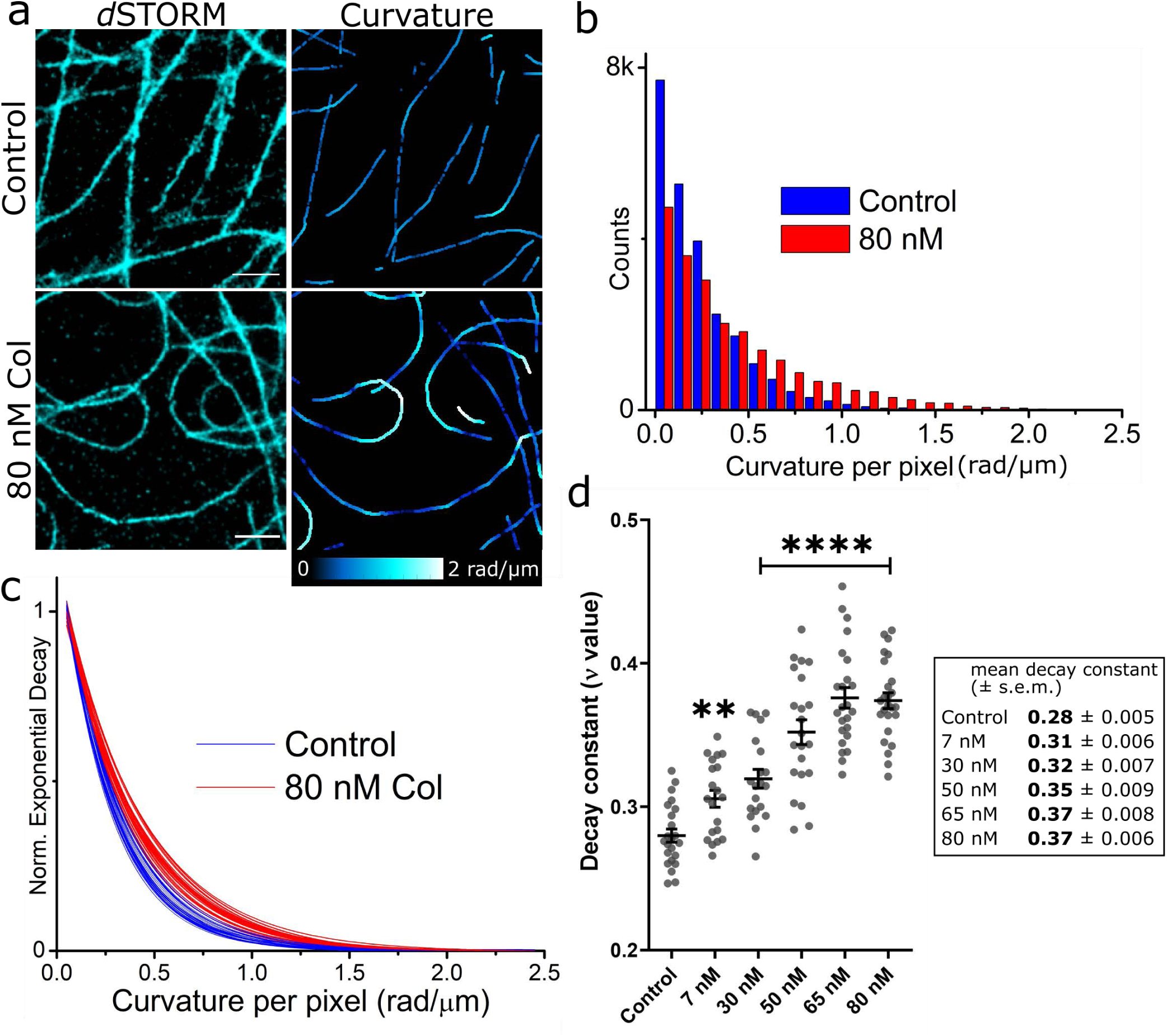
Analysis of filament curvature induced by colcemid. (a) Filaments from *d*STORM images of control and 80 nM colcemid treated HeLa cells (left). Curvature analysis using SIFNE and coloured at each traced pixel for curvature between 0 and 2 rad/*µ*m. SIFNE curvature output has been dilated by 10 pixels (using ImageJ) to enable visualization. Original SIFNE output shown in SI Figure 2a. Scale bars = 1 *µ*m. (b) Histogram of curvature data (curvature at each pixel) from a control cell and 80 nM colcemid cell with bins every 0.1 rad/*µ*m. Fitting of 80 nM cell data with residuals shown in SI Figure 2b. (c) Normalised exponential decays from histograms of control (N = 23) and 80 nM colcemid treated cells (N = 25). (d) Decay constants (*v* value) of each cell analysed with SIFNE (N = 135 cells, 50.4 mm total filament length) with mean and standard error of means. All *v* values were derived from exponential fits with adjusted R^2^>0.994. Parametric t-tests for each colcemid concentration against the control reveal significant difference with 7 nM colcemid (**, p = 0.0014) and each higher colcemid concentration 30 - 80 nM (****, p < 0.0001).

Curvature data histograms from each cell were fitted by a single exponential decay function in order to obtain a decay constant (presented as a reciprocal *v* value) which provides an empirical measure of the distribution of curvature where a larger *v* implies a greater proportion of higher curvature values in the histogram and therefore a larger ‘average curvature’ for the MT filaments of that cell. Figure 2c shows a distinct difference in the fitted exponential decay functions for control and 80 nM colcemid-treated cells. The same analysis was performed on all cells (0 - 80 nM) and the resulting *v* values are plotted in Figure 2d and show a clear trend of MT curvature increasing with colcemid concentration. Cells with the highest proportion of curvatures were those treated with 65 nM and 80 nM. With 100 nM and 200 nM colcemid treated cells however, we found SIFNE could not properly trace the few filaments present despite their prominence in *d*STORM images. The high density of speckled signal present in those cells interfered with SIFNE’s tracing algorithm to generate false filaments by joining specks together. These quantified results match the visual inspection of *d*STORM images where MTs are more curved but still intact at 65 nM and 80 nM, and clearly fragmented at higher concentrations.

Overall, *d*STORM paired with SIFNE can characterize both macro and sub-diffraction changes to MT curvature in response to colcemid treatments. Of the cells analysed for filament curvatures, the trend was that mean *v* values increased with increasing colcemid concentrations from 7 - 80 nM compared to the control. This suggests that filaments in the treated cells become more perturbed with additional colcemid, initially sustaining some mild curvature with lower doses (7 - 30 nM) then visually striking aberrant curvatures with higher doses (65 - 80 nM). Remarkably, even ultra-low doses were found to have significant effects on MT architecture despite no obvious filament perturbations observed in the corresponding *d*STORM images. Since these doses are associated with clinical use, mild filament curvatures can presumably be tolerated by cells in conjunction with therapeutic outcomes. The detection of MT structural effects, however mild, suggests that MT dynamics may also be impaired. Therefore we employed live-cell SOFI to probe MT filament dynamics in response to ultra-low colcemid doses.

### Sub-diffraction time-lapse imaging of live-cell microtubules correlates with fixed-cell imaging and reveals suppression of MT dynamics at ultra-low colcemid treatment

SOFI is advantageous for time-lapse imaging of live cells because it uses comparatively low excitation powers and can generate images in considerably less measurement time than *d*STORM. As a post-processing super-resolution technique, the degree of SOFI order applied determines the spatial resolution gain achievable (Figure 3). We compared 2^nd^ and 3^rd^ order for our data and while the resolution improvement is greater for 3^rd^ order, the better structural clarity produced with 2^nd^ order was chosen over the higher resolution of 3^rd^ order for tracing individual MT filament dynamics. Another parameter optimized was the duration of real-time raw data analyzed to render an individual super-resolution SOFI image (*I*_time_, see methods). The length of collated raw frames dictates not only image clarity, but also the overall temporal resolution when applied to time-lapse imaging. Because SOFI requires several seconds of acquired data to form one sub-diffraction image, any cell movement during this period will contribute blurry features to the SOFI image. The flexible motions of MT filaments in live cells can result in over-estimating widths or conceal multiple individual filaments as one wider filament. Although we found that *I*_time_ = 10 s was sufficient to generate single SOFI images we used *I*_time_ = 20 s for time-lapse movie generation in order to achieve higher clarity of individual MT filaments throughout the observation period.

**Figure 3.**
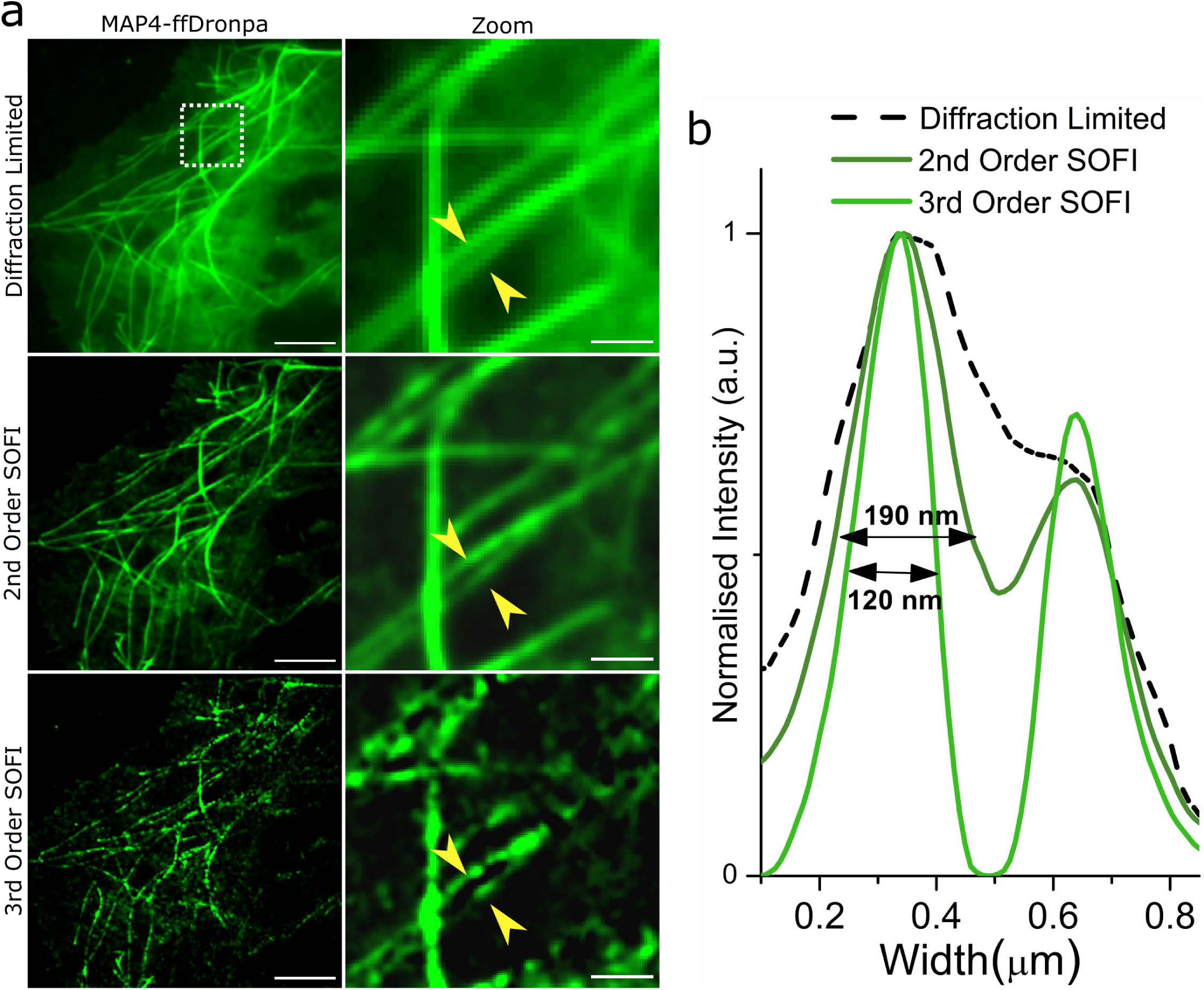
2^nd^ Order SOFI produces optimal trade-off between sub-diffraction resolution and image clarity of live-cell MT filaments. (a) Representative diffraction-limited, 2^nd^ and 3^rd^ Order SOFI images of MTs in a COS-7 cell labelled with MAP4-ffDronpa. Zoomed regions (dotted white box) show the same proximate filaments (yellow arrows) depicting between separation in the 2^nd^ and 3^rd^ Order renderings, but increased loss of structure and clarity in the 3^rd^ Order image. Scale bars = 5 *µ*m and 1 *µ*m. (b) Intensity cross-sections across the proximate filaments showing improved resolution with 2^nd^ and 3^rd^ Order SOFI. Values are FWHM of Gaussian fitted to the first peaks of the intensity plots. A Gaussian fitted to the first peak of the diffraction limited trace (dotted line) resulted in a FWHM of 450 nm, demonstrating the poor image resolution

From SOFI time-lapses, we found filaments could be clearly seen to grow and shrink, however, lateral motions as seen in Figure 4a were also observed and caused smearing in the SOFI image. Therefore, we focused on the growing and shrinking of filaments at the tips to provide a measure of MT dynamics. Figure 5a and 5b show sequential 20s SOFI images of representative filaments. Filament movement can be quantified by measuring growth and shrinking events based on just the filament tip (e.g. using a fluorescent label on MT tip protein EB3^44^), but because we were able to visualize a whole filament with sub-diffraction resolution, we measured filament growth and shrinkage by tracing several microns from a reference base to the filament tip. In this way, live MT activity was captured with SOFI and the changes in length (Δlength) of individual filaments over time quantified to elucidate MT dynamics with 20 second temporal resolution (Figure 5).

**Figure 4.**
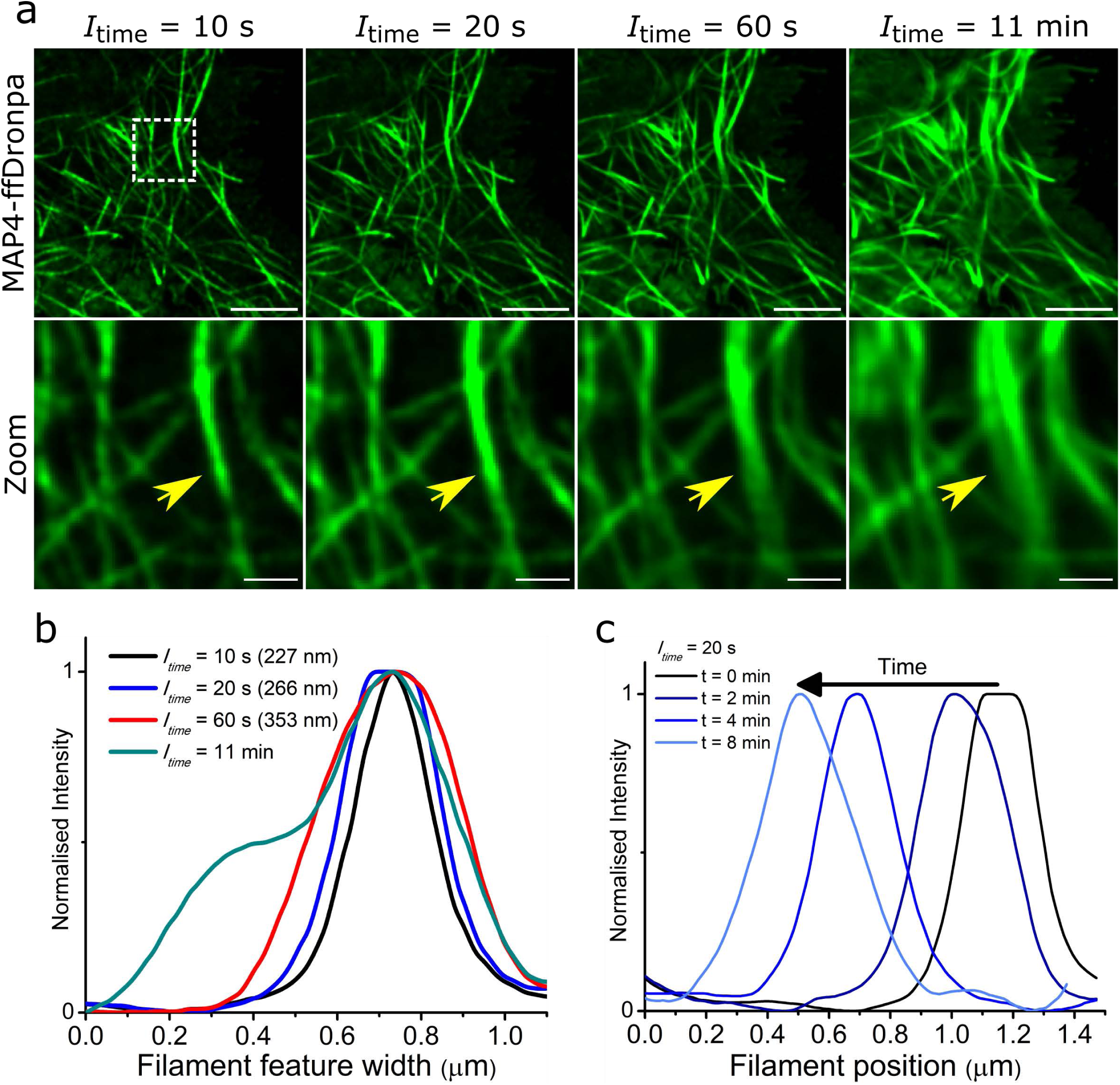
Shorter SOFI integration times reduce smear caused by filament movement while retaining sub-diffraction resolutions. (a) First frames from SOFI time lapse of live COS-7 MTs labelled with MAP4-ffDronpa and processed using different integration times (*I*_time_) from 10 seconds to 11 minutes. Zoom region (dotted white box) from each *I*_time_ image showing the same filament feature indicated by yellow arrow (possibly more than 1 filament) having increasing width with increasing *I*_time_. Scale bars = 5 *µ*m (top row) and 1 *µ*m (bottom row). (b) Intensity profile of filament feature width in (a) with different *I*_time_ and corresponding FWHM values after fitting with a Gaussian. (c) Filament feature width displacement through SOFI timelapse with *I*_time_ = 20 s at t = 0, 2, 4 and 8 min.

**Figure 5.**
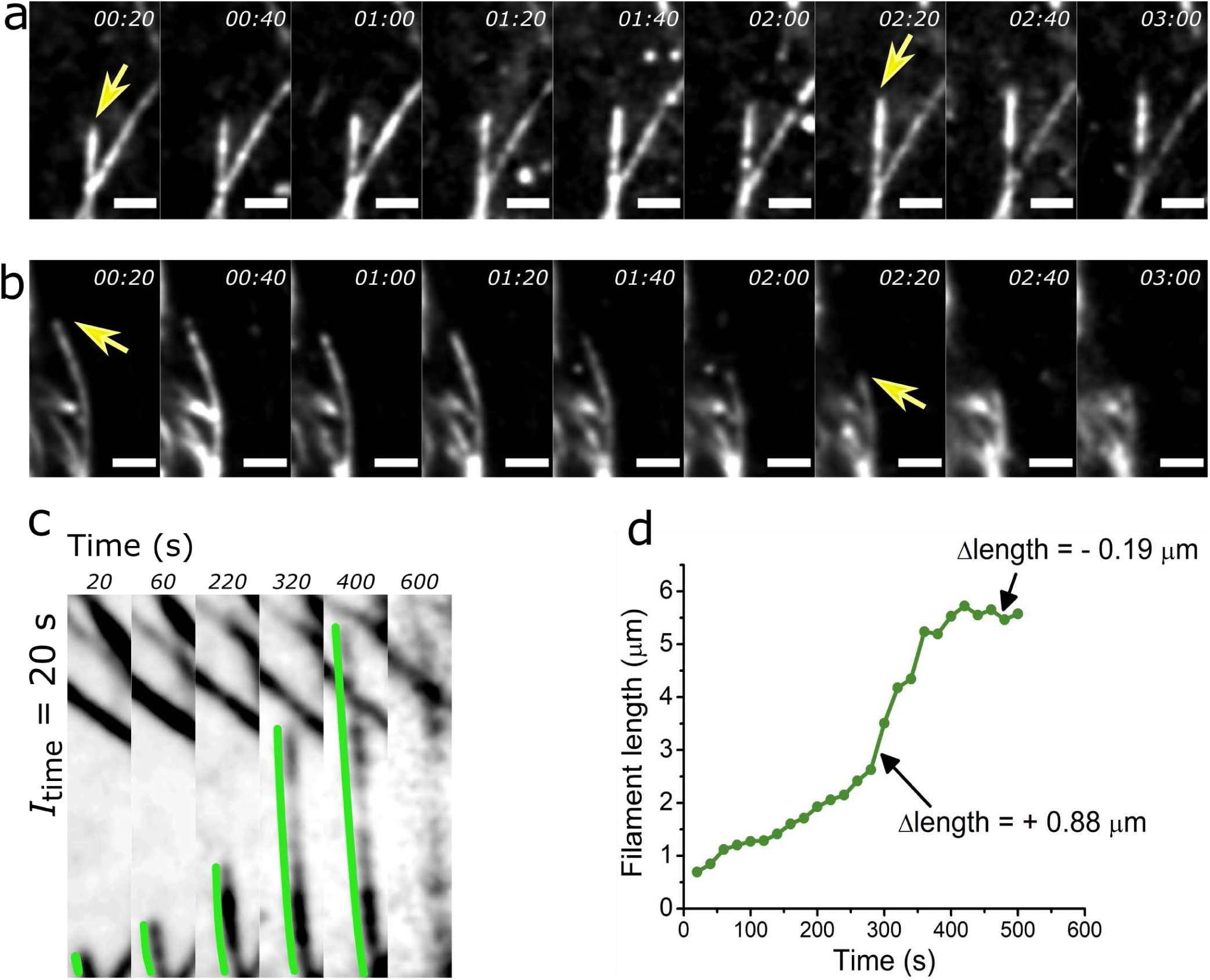
Tracing filament activity from SOFI time-lapse enables sub-diffraction changes in filament length to be quantified. (a-b) Consecutive SOFI frames of *I*_time_ = 20 s showing sub-minute dynamics of (a) growing and (b) shrinking MT activity in HeLa cells. Yellow arrows indicate filament of interest. Scale bars = 1 *µ*m. (c) Tracing a growing MT filament from a SOFI time-lapse. Green line measures length from reference base to tip, except in final frame of *I*_time_ = 20 s where filament becomes unclear. (d) Filament from (c) traced at each frame for ∼10 minutes and example measurements for both growth and shrinkage Δlength between successive frames.

Static SOFI images of colcemid treated cells show similarities to those imaged with *d*STORM (Figure 6a). Cells treated with 7 - 50 nM colcemid had filaments of similar appearance to the control cells with mostly linear filaments extending outward from the centre. At 65 nM and 80 nM colcemid, there was more curvature present with more filaments curved inward away from the cell edge. In 100 nM colcemid treated samples a small number of abnormally short filaments could be detected. SIFNE, however, could not accurately trace individual filaments for curvature analysis given the high MT density in 2^nd^ order SOFI images. Overall, the trend from visual inspection of live cells matches that observed from *d*STORM and therefore demonstrates that abnormal curvatures observed and quantified using *d*STORM were not a result of fixation artifacts. We also hypothesize that the observable increase in unstructured ‘non-specific’ fluorescence signal seen in ≥65 nM colcemid-treated cells corresponds to the well-resolved speckled signals detected in the *d*STORM images. This is explained by the increase in MT breakage and disassembly resulting in higher concentrations of dimeric/oligomeric tubulin. While these individual species could be resolved as speckles in the *d*STORM images of fixed cells, the poorer resolution achieved with 2^nd^ order SOFI, combined with the dynamic movement of these species within a live cell resulted in a higher intensity of unstructured blur. The effects of these dynamics combined with out-of-plane fluorescence also manifest as the uneven filament signal observed in cells imaged with SOFI (SI Figure 3). Because correlation is based on the quality of fluctuations - weaker fluctuations further away from the focal plane (higher planes) will yield dimmer SOFI-calculated pixels meaning those MTs perfectly in focus present more brightly than those slightly out of focus.

**Figure 6.**
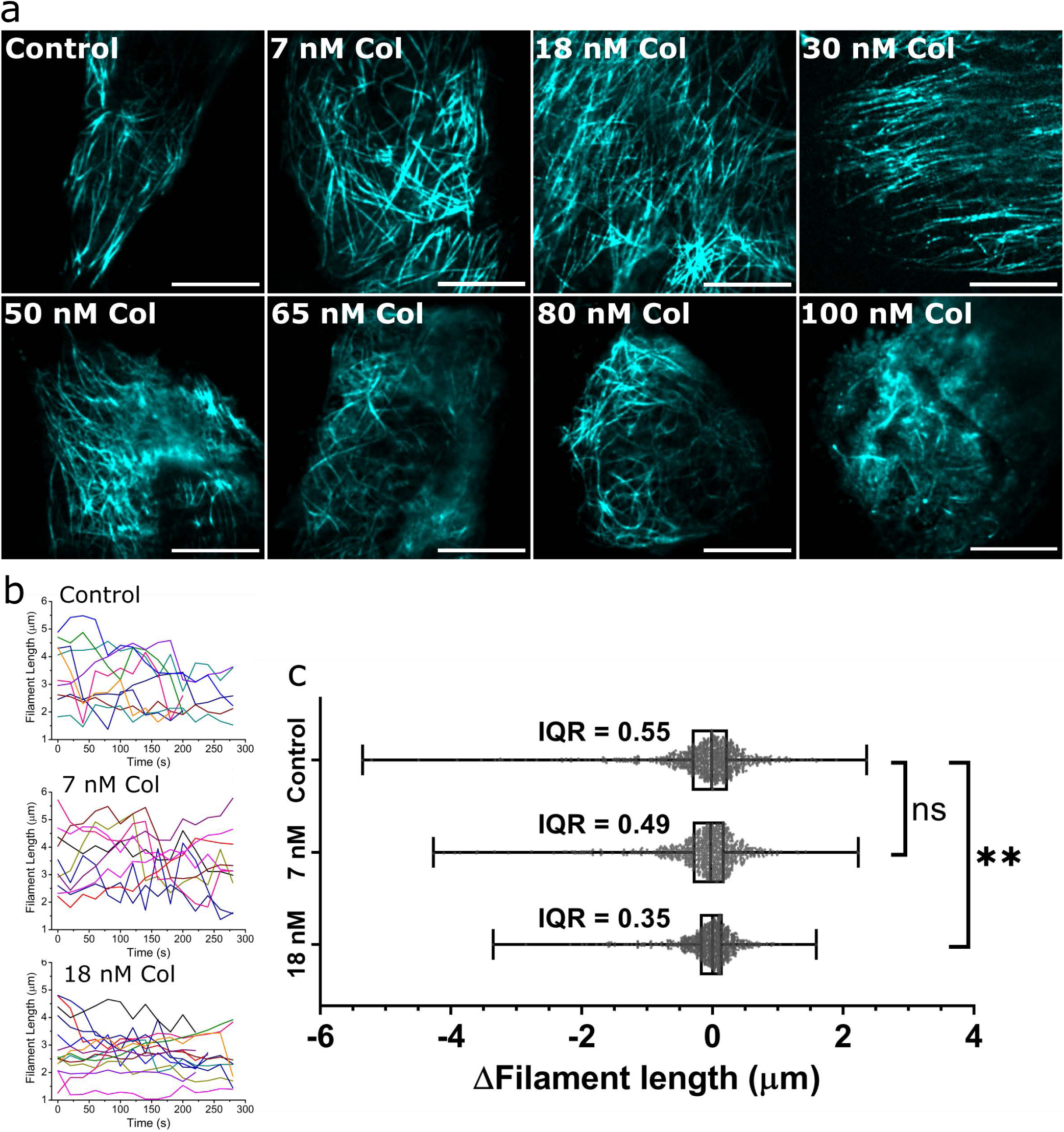
SOFI reveals colcemid causes filament curvature and suppresses filament dynamics in live cells. (a) SOFI of live HeLa MTs labelled with MAP4-ffDronpa, imaged after 5 hours treatment with the specific concentration of colcemid. Live-cell MTs imaged with SOFI show similar trend to fixed-cell MTs imaged with *d*STORM. Aberrant filament curvature were observed in 65 nM and 80 nM colcemid treated cells, then some degree of fragmentation became evident in 100 nM colcemid treated cells. Each cell is representative of each condition imaged with SOFI (SI Figure 3). Scale bars = 5 *µ*m (b) Overlay of 10 representative filament traces from SOFI time-lapses of control, 7 nM and 18 nM colcemid treated cells. Representative SOFI time-lapse movies of each condition is shown in SI movie 1. (c) Length values were extracted from between each consecutive frame and used to determine the given Δlength for each pair of frames. Distributions of Δlength values obtained from 3 independent assays, N = 12 cells, >96 filaments and >770 Δlength events for each condition. Scatter plots of data overlaid with median, interquartile range (IQR, box) and full range (whiskers). Kolmogorov-Smirnov test used to determine significant difference between control and 18 nM colcemid treatment (**, p = 0.0011).

For SOFI time-lapse assays, we applied 2^nd^ order correlation to achieve both sub-diffraction resolution and high image clarity of individual live-cell MT filaments. We also used *I*_time_ = 20 s for the best temporal resolution that retained the clarity of filaments while minimizing the amount of smearing caused by lateral filament movements. By tracing the Δlength changes of individual filaments over several minutes, we were able to quantify instants of both growth and shrinkage. These optimized SOFI parameters were used to interrogate ultra-low colcemid concentrations (7 nM and 18 nM) where although no visible filament deformations were observed in either the *d*STORM or SOFI images, significant increases to filament curvature hinted that MT dynamics could also be affected. Because MT filaments behave differently depending on their position in the cell^45^, sampling of filaments for dynamics imaging and tracing was consistent i.e. all measured filaments were from the edges of cells with the presumption that colcemid effects would be most prominent here. Figure 6b shows traces of 10 representative filaments from each condition, where 18 nM colcemid treated filaments had relatively smoother traces than the control cell traces, indicative of slower dynamics. Extracting Δlength provided multiple measurement points (14 Δlength values for a 5 minute time-lapse) from a single filament which could be compiled and visualized as the total distribution of Δlength for each condition. This analysis revealed that the overall distribution of dynamics in control and 18 nM colcemid treated cells was significantly different with the colcemid treatment causing a loss of overall dynamic movement with both growth and shrinkage occurring more slowly (Figure 6d). These data demonstrate a loss of MT dynamics at ultra-low colcemid concentrations and reveals previously uncharacterised nanoscale dysfunction corresponding to therapeutic dosage regimens.

## Discussion

The observed responses of MTs in the presence of colcemid include suppressed dynamics, filament curvature and filament fragmentation (Figure 7). These MT alterations provide insight into drug mechanisms at the subcellular scale, but are also useful for understanding clinical outcomes. Colchicine (and colcemid) binds free tubulin subunits^11^ to form tubulin-colchicine complexes that hinder further filament growth (inhibits further tubulin polymerization) when incorporated into growing MT filaments^46^. Using SOFI time-lapses, we observed this effect where 18 nM colcemid suppressed filament growth but interestingly also reduced the events of filament shrinkage, implying gained filament stability and/or slowed rate of disassembly. With overall dynamics suppressed, MTs would be less able to form mitotic apparatus and generate sufficient pulling force to segregate chromosomes. Since this concentration falls within the range of clinically therapeutic doses, suppressed dynamics could be a means to prevent tumours from spreading by inhibiting cancer cell migration due to restricted motility. A previous study showed a similar range of colchicine (12 nM and 25 nM) significantly reduced the ability of human cells to migrate through 8 *µ*m plastic pores^47^. The lack of aberrant filament curvatures observed up to 30 nM suggests MT-dependent intracellular transport may still remain viable, such that these lower concentrations are relatively less harmful for non-cancer cells. Measuring the dynamics of other components such as motor proteins could reveal if trafficking/signalling are similarly suppressed by these clinically acceptable levels of MT-interacting drugs.

**Figure 7.**
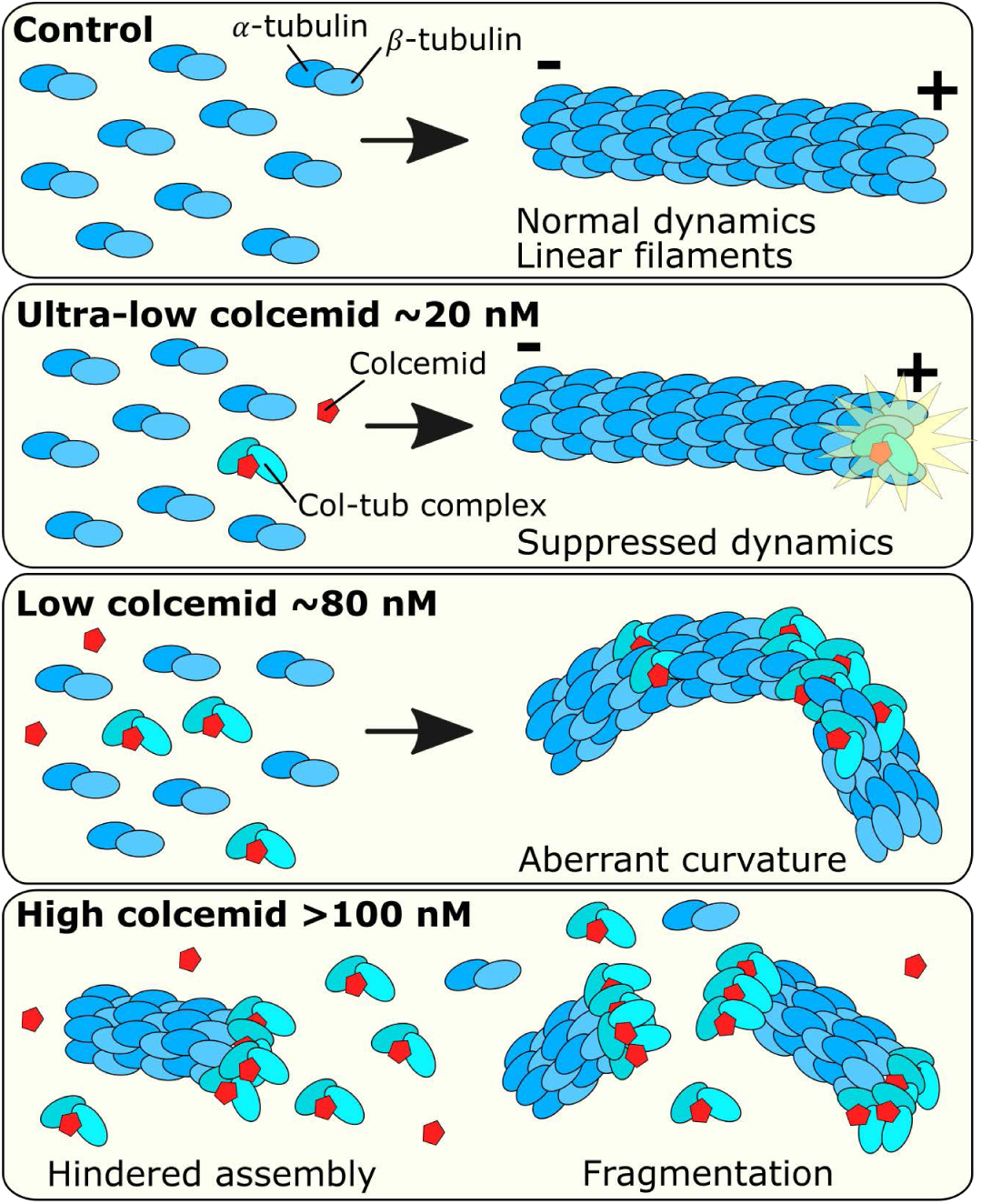
Microtubule dysfunctions with different doses of colcemid. MT filaments are polymers of *α*- and *β* -tubulin subunits that assemble through dynamic processes. Filaments grow toward a positive end and form relatively linear filaments that extend outward from the cell centre toward the cell membrane. With ultra-low levels of colcemid, MT dynamics are suppressed and filaments grow and shrink at a slower rate. Additional colcemid added produces more tubulin-colcemid complexes that distort normal tubulin-dimer subunit configurations, resulting in aberrant filament curvatures when these abnormal complexes become incorporated. High levels of colcemid produce filament fragments, either due to curvature strains that result in breakage or the inability for filaments to form properly due to an excess of tubulin-colcemid complexes.

The binding of colcemid at the colchicine binding site in between *α*- and *β* -tubulin monomers distorts the protein configuration of tubulin subunits^48^. The resulting misshaped *αβ* -tubulin-colcemid complex interferes with normal end-to-end assembly of typically linear MT filaments, introducing a kink into the filament and subunits become oriented at various angles to one another. With enough of these, filament curvatures become more apparent. This is consistent with the increasing frequency and extent of curvatures observed with increasing colcemid concentrations, starting from as low as 50 nM, followed by the appearance of curvatures beyond 2 rad/*µ*m at 65 nM and 80 nM. Such drastic remodelling of MT architecture is bound to impact the efficiency of intracellular transport and signalling, but since 80 nM is an upper limit of therapeutic dose (for colchicine), cells may be able to survive with some aberrant MT curvature. At 100 nM and 200 nM some curved MTs are observed together with shorter filament fragments, suggesting that the fragmentation may occur as a result of excessive structural strain at these highly curved sites. Also, since these higher concentrations result in a larger proportion of drug-affected tubulin in cells, MT filaments may find it more difficult to sustain assembly using native tubulin. Therefore, subunits are less likely to be incorporated into filaments but remain as free cytoplasmic constituents (dimers/oligomers), supported by the observation of speckled features in *d*STORM imaged cells after high colcemid treatments. Inducing MT filament fragments, either through curvature-derived breakage or inhibition of tubulin assembly, reflects a toxic outcome (≥100 nM) because most the MT network is absent. The direct MT architecture perturbations observed would likely have wide-ranging consequences for various vital MT functions including intracellular transport, cellular mobility and tissue structure. Additionally, reduced filament dynamics would likely affect genetic segregation during mitosis even if the overall structure of the spindle could be formed. Furthermore, our studies highlight contemporary challenges and limitations associated with characterization of nanoscale cellular landscapes, particularly more subtle drug effects that are likely of high importance in therapeutic understanding.

In conclusion, we have applied complementary super-resolution imaging methodologies to study the impact of colcemid on the dynamics and structures of MTs. We find that increasing colcemid concentration correlates with increasingly distinct MT effects in HeLa cells where the transition between each observed effect is sensitive to concentration increments of several tens of nanomoles. Interestingly, the three MT effects coincide with colchicine at normal therapeutic concentration in blood (18 nM: suppressed MT dynamics), maximum therapeutic concentration in blood (80 nM: MT curvature), and toxic concentration in blood (100 nM: MT network loss)^36^. From this, we suggest filament curvature to be a visually striking marker for impending toxicity of colcemid and possibly other colchicine derivatives. The onset of these MT perturbations after only a short exposure to the drug support the notion that traditional ‘antimitotics’ affect MT function even outside mitosis. There remain many challenges to directly observe and quantify dynamics of continuous fluid-like structures such as MT filaments, no less in super-resolution or under the influence of drugs. Improving both spatial and temporal resolutions will provide closer to realistic visualization of intracellular components and how they respond to clinical therapies.

## Methods

### Cell culture

HeLa (human epithelial adenocarcinoma, ATCC CCL-2) and COS-7 (african green monkey kidney fibroblast-like, ATCC CRL1651) cells were cultured in Dulbecco’s Modified Eagles Medium (DMEM – high glucose) supplemented with 10% fetal bovine serum and 1% penicillin-streptomycin, and incubated at 37°C with 5% CO_2_. Cell stocks were passaged twice a week to maintain 40 - 90% confluence in 25 cm^2^ flasks. For live-cell imaging, cells were seeded onto high precision coverglass (Marienfeld 18 mm diameter #1.5H coverglasses, cat #0117580), and grown to 60% confluence before transfection. Seeded cells were transfected using Fugene HD Transfection Kit according to manufacturer’s instructions (Promega). For each chamber, transfection media consisted 500 ng of DNA plasmid (pMAP4-N1-ffDronpa) and 2 *µ*L Fugene reagent in 50 *µ*L DMEM. Cells were incubated in tranfection media for at least 18 hours. For drug assays, colcemid (Roche cat #10295892001) was added to cell growth media to yield final concentrations ranging from 7 to 200 nM 5 hours before fixation and immunostaining, based on a previously optimised protocol^49^. Typically, colcemid ranging 100 – 300 nM (∼40 - 100 ng/ml) is used in laboratory settings to induce mitotic arrest for synchronizing cell cultures or for chromosome spreading protocols^35^.

### *d*STORM sample preparation

For *d*STORM imaging, untransfected Hela cells were cultured and treated with colcemid as above before being pre-permeabilised in 0.25% Triton X-100, 0.3% glutaraldehyde (Alfa Aesar cat #A17876) in cytoskeletal buffer (CB: 10 mM MES pH 6.1, 150 mM NaCl, 5 mM EGTA, 5 mM MgCl_2_, 5 mM Glucose) at 37°C for 30 seconds, then fixed in 2% glutaraldehyde in CB for 10 minutes at 37°C. Fixed cells were washed in phosphate buffered saline (PBS) twice for 5 minutes, then quenched in 0.1% NaBH_4_ in PBS for 7 minutes at R.T, then washed twice in PBS for 5 minutes. Cells were blocked in 5% bovine serum albumin in PBS for at least 30 minutes at R.T. before immunostaining with rabbit anti-*α*-tubulin (Abcam ab18251, 1:500 in 5% BSA/PBS, 1 hour, R.T) then anti-rabbit Alexa Fluor 647 conjugate (ThermoFisher cat #A-21246, 1:200 in 5% BSA/PBS, 45 min, R.T). Following each antibody stain, cells were washed in 0.1% Tween-20 in PBS twice for 5 minutes. Cells were then post-fixed in 3.7% formaldehyde for 5 minutes at R.T. A switching buffer of 100 mM mercaptoethylamine (MEA) in PBS made to pH 8.2 (adjusted with KOH) was added to cells for *d*STORM imaging.

### Single molecule super-resolution imaging

Imaging was performed on a home-built single molecule super-resolution widefield microscope as previously described^50^. Briefly, we used an Olympus IX81 inverted fluorescence microscope frame fitted with a TIRF 100X 1.49 NA oil objective, Oxxius 638-nm and Toptica 488-nm laser diodes, and Andor iXon EM-CCD detector. Acquisition parameters were controlled using Micromanager. For *d*STORM, cells were imaged in a switching buffer of 100 mM mercaptoethylamine (MEA) at pH 8.5 in PBS. The 638-nm laser was used at full power (150 mW resulting in 3 - 5 kW/cm^2^) to induce photoswitching of Alexa Fluor 647 for resolved single molecule emissions (observed as “blinking” events) that were acquired using Micromanager^51^ (20 ms exposure, 100 gain, > 10,000 frames). Imaging was performed in quasi-TIRF mode that captured emissions from a limited axial range of a few micrometers above the coverslip. Acquired frames were analysed in rapi*d*STORM^52^ using input pixel size 100 nm and point spread function full width half maximum (PSF FWHM) of 360 nm to localise each single molecule emission. A reconstructed coordinate map of all localisations (over 1 million per image) produced the 2D super-resolved *d*STORM image.

### Filament curvature analysis

The contrast of *d*STORM images were enhanced with a Gaussian blur (ImageJ^53^, sigma (radius) = 1) to enable better MT tracing from SIFNE^43^. For measuring filament curvature, MT images were processed through SIFNE with default parameters except “Max Curvature” which was set to 3 rad/*µ*m. This was to accommodate detection of more extreme curvatures and curvatures occurring along the axial plane (curvature along the z-axis would be more pronounced when visualised from orthogonal 2D perspectives). At the last step of the SIFNE analysis to avoid potentially fragment filaments, a minimum length of 200 nm was set for inclusion in the analysis. Curvature of each pixel traced by SIFNE (at least 14,000 per cell) were compiled into a histogram (0.1 rad/*µ*m bins) and fitted with a single exponential decay function:

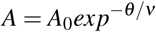

where A is the proportion of curvature events with a given curvature *θ*, A_0_ is the initial amplitude at *θ* = 0, *v* is the reciprocal of the first order decay rate constant. Cells from 2 independent assays of each colcemid condition were compiled to determine the mean decay constant. Unpaired parametric t-test with Welch’s correction were performed on compiled *v* values for each colcemid concentration against the control. Significant difference was found with 7 nM colcemid treatment (**, p = 0.0014) and each higher colcemid treatment 30 - 80 nM (****, p < 0.0001).

### SOFI acquisition

SOFI was performed on the same setup as *d*STORM experiments. HeLa cells were grown on coverglasses and labelled by transfection with a DNA plasmid for transient expression of ffDronpa conjugated to microtubule associating protein 4 (MAP4). Just before imaging, transfected cells were rinsed with warm PBS then mounted in a custom-built chamber and filled with warm PBS. Upon excitation with continuous 488-nm laser at ∼50 mW/cm^2^ (Toptica, total output = 2 mW), ffDronpa photoswitched at rates suitable for SOFI analysis. ffDronpa was also stably fluorescent at even lower power (∼5 mW/cm^2^, 0.2 mW), however was found to be insufficient for photoswitching. Ideal raw data for SOFI is a dense coverage of fluctuations across the labelled structure. As such, the general shape of MT filaments is clearly visible during SOFI acquisition, unlike for *d*STORM where only single molecule emissions are observed. Relevant to both techniques is that the quality of raw data determines the quality of super-resolution images. During acquisition, cells were exposed to cumulative laser exposure for no longer than 10 minutes. Live-cell SOFI raw data acquisition was performed using Micromanager with 100 gain at 20 Hz, and saved as .tif movie stacks. Between 2,000 – 6,000 frames were collected from each cell depending on signal quality over time.

### SOFI processing

Acquired frames were imported into the Localizer package^54^ in Igor Pro 7. Under the SOFI tab, the following parameters were applied for static SOFI images; Order = 2, Pixel Combos = More, Also average image = checked, Frames: 0-399. Executing the analysis produced: i) the average image combining all acquired frames, representative of a diffraction-limited fluorescence image; ii) the correlated SOFI image. Finally, a Richardson-Lucy deconvolution was applied; Standard deviation of the PSF = 1.6 pixels, Number of iterations = 2. The contrast of the image was further modified by selecting Macros>Append colourscale sliders and adjusting upper and lower limits of the final SOFI image. The processing sequence is identical to create a SOFI time-lapse, with the exception of selecting “Make movie” and selecting the number of acquired frames to be correlated into each SOFI frame. Acquired data of several thousand frames were correlated every 400 frames (accounts for 20s of real time data) to produce a continuous SOFI time-lapse up to 5 minutes. Integration time (*I*_time_) describes the duration of real time data used to generate one SOFI image or one frame in a SOFI time-lapse (Figure 4). The Localizer^54^ interface enables control of *I*_time_ by selecting the number of acquired frames per SOFI image: 200 frames = 10 s, 400 frames = 20 s, 1,200 frames = 1 min, 13,000 frames = 11 min. Lower integration times produced more fluid SOFI time-lapses of MT dynamics (higher temporal resolution), but lacked improved spatial resolutions and image clarity of higher integration times.

### SOFI orders for improving resolution and clarity

To test correlation parameters, the same data were processed using 2^nd^ and 3^rd^ order SOFI correlations (Figure 3a). Compared to the diffraction-limited fluorescence image, both SOFI correlation orders enhanced the contrast and clarity of individual MT filaments. In zoomed in regions, it becomes clear that the number of pixels increases (and pixel size decreases) through the formation of ‘virtual pixels’ inherent to the SOFI process^55^. Using FWHM of filament intensity cross-sections, widths were found to be in the range 120 – 190 nm, which is about a factor of 2 to 3 improvement over the measurement from the original diffraction limited image (unresolved ∼450 nm) as expected for 2^nd^ and 3^rd^ order SOFI analysis respectively. Although this is not as good a resolution gain as can be achieved using *d*STORM, SOFI provides more biological relevance with live-cell imaging. Comparing between SOFI orders, we found 3^rd^ order yielded better distinction of two adjacent filaments than 2^nd^ order. However, a significant amount of filaments correlated by 3^rd^ order were discontinuous and the overall MT network was less visible. Despite the additional resolution improvement from 3^rd^ order, we used 2^nd^ order SOFI for all subsequent analysis, given its consistency in retaining the integrity of whole filaments throughout the entire image. Subsequent SOFI for static images and time-lapses were performed using consistent acquisition parameters (20Hz acquisition framerate for at least 20s) and processed with 2^nd^ order SOFI correlations.

### Optimizing *I* _time_ for SOFI time-lapse

To achieve clear MT filaments throughout the time-lapse, we required a minimum *I*_time_ of 20 s (400 acquired frames acquired at 20 Hz) per SOFI frame which corresponds to 20 s temporal resolution. We tested different integration times (*I*_time_ = 10 s, 20 s, 1 min, 11 min) on acquired data of living MTs to make SOFI time-lapses with different temporal resolutions (Figure 4a). The movement of MTs throughout acquisition, when integrated, produced smearing artefacts with longer *I*_time_ that added apparent size to structures. A measured filament feature (possibly more than one filament) increased in width from 230 nm up to >500 nm by increasing *I*_time_ from 10s to 11 min (Figure 4b) due to its lateral movement during acquisition. Using *I*_time_ = 20 s, we could track its displacement across several hundred nanometers after 2, 4 and 8 min of real time with a similar width retained throughout (Figure 4c). Though filaments could be rendered using *I*_time_ = 10 s for better temporal resolution, each SOFI frame had relatively less signal and it was more difficult to resolve continuous structures. This was especially apparent after ∼9 minutes of continuous imaging where ffDronpa signal was reduced due to photobleaching. By using *I*_time_ = 20 s for all SOFI time-lapse experiments (and previously shown SOFI images), we compromise temporal resolution for longer observation times of MT dynamics with better filament contrast and clarity.

### Tracing live MT filaments

Filaments at the cell edge were selected for analysing dynamics because colcemid-induced depolymerization occurs from filament (+) ends, and so we hypothesized that initial interruption to MT dynamics would manifest most prominently at the edges. Additionally, cells are typically thinnest at the edges compared to the centre, meaning imaging at these areas minimized the amount of out-of-focus fluorescence, improving the quality of the raw and rendered SOFI images. We used the segmented line tool in ImageJ to trace each filament by selecting a reference base point and measured to the tip at each SOFI frame throughout the SOFI time-lapse (Figure 5c). Each length was plotted as a function of time to form a filament trace (Figure 5d). Because photobleaching of RSFPs becomes more apparent in the later acquisition frames labelled structures in later SOFI time-lapse frames may not be as clear. The example here is the filament (imaged with *I*_time_ = 20 s) at t = 600 s 5c that is barely visible and could not be traced because of the degrading RSFP signal. However, at least 5 minutes of traceable filaments was obtainable from each cell and up to 9 minutes in some instances.

### Measuring drug-affected microtubule dynamics

We performed triplicate assays of each colcemid treatment (0 nM, 7 nM, 18 nM) for measuring microtubule dynamics using SOFI. We sampled at least 8 filaments from each cell (4 cells per assay) using SOFI time-lapse movies. In each SOFI frame, filaments were measured from a reference base to the tip using the segmented line tool in ImageJ. Measurements were used to plot change in length (Δlength) of each filament every 20 s. All Δlength (+ve and –ve) were compiled for each drug condition as a scatter plot and overlaid with a box-and-whisker plot of the interquartile range (IQR) and full range (whiskers). Δlength values when combined were not normally distributed, failing normality tests (D’Agostino-Pearson test and Shapiro-Wilk test in GraphPad Prism 8). This is consistent with the idea that accumulated MT growth and shrinkage rates would not have normal distributions given the presence of other proteins that regulate MT dynamics^56^ and that the rate of MT disassembly is typically faster than assembly. Given the non-normal distribution, we used the IQR to describe the spread of Δlength values. Compared to the control, we observed a narrower distribution from the 18 nM colcemid data, indicative of suppressed MT dynamicity i.e. the lengths of traced filaments increased and decreased less or at slower rates during the periods of observation. The reduced IQR shows overall hindered filament activity by about 11% and 36% with 7 nM and 18 nM colcemid treatment respectively. Applying the Kolmogorov-Smirnov test, we found 18 nM colcemid induced a significant difference (**, p = 0.0011) to the distribution of Δlength values compared to the control.

## Acknowledgements

Support from the Australian Research Council through its Discovery program (DP170104477) is gratefully acknowledged. Dr Whelan is the recipient of an Australian Research Council Australian Discovery Early Career Research Award (DE200100584) funded by the Australian Government.

## Author contributions statement

AMR, DRW and TDMB conceived the experiments. AMR prepared and imaged the fixed cell *d*STORM samples. AMR, SD and RBH prepared live cell samples and performed SOFI imaging. AMR and CE analysed the results. AMR wrote the manuscript with all authors editing and reviewing the manuscript.

## Additional information

The authors declare no conflicts of interest.

